# Atlantean Evolution in Darwin’s Finches—Issues and Perspectives in Species Delimitation using Phenotypic Data

**DOI:** 10.1101/124610

**Authors:** Carlos Daniel Cadena, Iván Jiménez, Felipe Zapata

## Abstract

Progress in the development and use of methods for species delimitation employing phenotypic data lags behind conceptual and practical advances in molecular genetic approaches. The basic evolutionary model underlying the use of phenotypic data to delimit species assumes random mating and quantitative polygenic traits, so that phenotypic distributions within a species should be approximately normal for individuals of the same sex and age. Accordingly, two or more distinct normal distributions of phenotypic traits suggest the existence of multiple species. In light of this model, we show that analytical approaches employed in taxonomic studies using phenotypic data are often compromised by three issues: (1) reliance on graphical analyses of phenotypic space that do not consider the frequency of phenotypes; (2)exclusion of characters potentially important for species delimitation following reduction of data dimensionality; and (3) use of measures of central tendencies to evaluate phenotypic distinctiveness. We outline approaches to overcome these issues based on statistical developments related to normal mixture models and illustrate them empirically with a reanalysis of morphological data recently used to claim that there are no morphologically distinct species of Darwin’s ground-finches (*Geospiza*). We found negligible support for this claim relative to taxonomic hypotheses recognizing multiple species. Although species limits among ground-finches merit further assessments using additional sources of information, our results bear implications for other areas of inquiry including speciation research: because ground-finches have likely speciated and are not trapped in a process of “Sisyphean” evolution as recently argued, they remain useful models to understand the evolutionary forces involved in speciation. Our work underscores the importance of statistical approaches grounded on appropriate evolutionary models for species delimitation. Approaches allowing one to fit normal mixture models without *a priori* information about species limits offer new perspectives in the kind of inferences available to systematists, with significant repercusions on ideas about the structure of biological biodiversity. [morphology; normal mixture model; phenotype; principal components analysis; species limits; variable selection.]

---

Systematic biology seeks to discover and describe species, and to establish phylogenetic relationships among them and among clades at higher levels. Given these two main goals of the field, reviews published over a decade ago noted that the literature on theory and methods of phylogenetic inference and on theory of species concepts was extensive, whereas methods for delimiting species had received much less attention (Sites and Marshall 2003; Sites and Marshall 2004). Over the past few years, this imbalance has been partly overcome with considerable development, application, and integration of methods for species delimitation (Padial et al. 2010; Camargo and Sites 2013). Largely driven by increased availability of multilocus datasets brought about by advances in DNA sequencing technology, however, much recent progress has focused on probabilistic methods for analyses of molecular data (reviewed by Fujita et al. 2012; Carstens et al. 2013), whereas relatively little effort has been devoted to approaches using phenotypic data to delimit species (Wiens and Servedio 2000; Ezard et al. 2010; Guillot et al. 2012; Zapata and Jiménez 2012; Edwards and Knowles 2014; Solís-Lemus et al. 2014). Yet, because most fossil and living species have been discovered and named based on phenotypic distinctiveness (Luckow 1995; Mallet 2013; Miller 2016), and because genomic-based species delimitation approaches are no substitutes for judicious assessments of other sources of information (Sukumaran and Knowles 2017), the theory and practice of delimiting species using phenotypic data remain central to modern systematics.

Although species descriptions employing phenotypic data are often non-quantitative and although systematists may often not be explicit about the rationale they follow to delimit species (Luckow 1995; McDade 1995; Sangster 2014; Allmon 2016), the use of objective criteria for species diagnosis based on phenotypic characters has a long tradition in taxonomy, rooted on evolutionary theory (Wiens and Servedio 2000; Zapata and Jiménez 2012; Futuyma 2013). The basic evolutionary model for the distribution of a continuous quantitative character within a species (Fisher 1918) assumes polygenic inheritance and random mating; under these assumptions, gene frequencies are expected to be close to Hardy-Weinberg equilibrium and phenotypic variation among individuals of a single species tends to be normally distributed (Templeton 2006). On the other hand, if phenotypic variation is best described by two or more distinct normal distributions, then one may conclude that there is more than one species in a sample of individuals (Coyne and Orr 2004; Mallet 2008). This conclusion is granted under the assumption that distinct normal distributions in polygenic traits do not reflect age- or sex-related variation, or phenotypic plasticity. It follows that distinct normal distributions in cases in which phenotypic variation is caused by few loci of large effect (e.g., Smith 1993) or largely driven by environmental factors (e.g., Moczek and Emlen 1999) do not constitute evidence of more than one species. Finally, because distinct phenotypic distributions may represent evidence of species boundaries given a variety of criteria for species delimitation (Luckow 1995; Zapata and Jiménez 2012), the Fisherian model described above serves as a conceptual basis to establish species limits under multiple species definitions (sensu de Queiroz 1998).

Despite the long tradition of this basic model for species delimitation based on quantitative phenotypic characters, statistical tools for its formal application to empirical data were limited until recently. Procedures allowing one to fit combinations of normal distributions to phenotypic variation among specimens, without *a priori* knowledge of species limits, were initially developed in the late XIX century (Pearson 1894). However, practical application only became possible following computational advances in the 1970s (i.e., the expectation–maximization algorithm; McLachlan and Peel 2000) and software development from the late XX century into the present (Fraley and Raftery 2002; Fraley et al. 2012). Because these statistical approaches entered the literature on species delimitation only a few years ago (Ezard et al. 2010), it is not surprising that even recent studies do not employ them when analyzing phenotypic data to delimit species. Instead, systematists frequently infer species limits examining phenotypic variation based on visual inspection of scatter plots defined by a few axes that account for most phenotypic variance, often derived from principal components analysis (PCA). In addition, systematists often delimit species based on differences between groups of specimens in the central tendency of phenotypes. This is true of work on living plants and animals (reviewed by Rieseberg et al. 2006), as well as in studies of extinct taxa in the fossil record (reviewed by Allmon 2016).

Here, we show that analytical approaches commonly employed in taxonomic studies are inadequate in light of the evolutionary model underlying species delimitation described above. It follows that if species delimited by inadequate statistical approaches are used as units for subsequent analyses, then any mistakes may carry on and influence views in other areas of inquiry, such as speciation research. Focusing on Darwin’s finches from the Galapagos Islands, an iconic group for the study of natural selection, speciation, and adaptive radiation (Lack 1947; Bowman 1961; Grant 1999; Grant and Grant 2008; Grant and Grant 2014), we provide an example of how employing statistical approaches explicitly related to the basic evolutionary model underlying the use of phenotypic data in species delimitation may enhance assessments of species limits and thus our understanding of evolutionary processes.

## Sisyphean Evolution in Darwin’S Finches?

Among Darwin’s finches, the many studies of ground-finches in the genus *Geospiza* have been especially productive in terms of insights into species formation and the role of geographic isolation, natural selection, and hybridization in microevolutionary processes that may scale up to macroevolutionary patterns (reviewed by Grant 1999; Grant and Grant 2008; Grant and Grant 2014). There has been considerable disagreement in the literature about the number of species in the group (reviewed by McKay and Zink 2015), but most modern taxonomic treatments have recognized six species of ground-finches (Lack 1947; Rising et al. 2011). However, based on genomic evidence (Lamichhaney et al. 2015) and some vocal and behavioral data, three subspecies were recently elevated to species rank, bringing the total number of recognized species to nine (Remsen et al. 2017).

In a provocative recent paper, however, McKay and Zink (2015) offered an intriguing alternative perspective on the taxonomy and evolution of ground-finches (see also Zink 2002). These authors boldly argued that morphological evidence for the existence of multiple species of *Geospiza* is lacking, and they presented the iconoclastic argument that different phenotypes should be considered transient ecomorphs within a single species. Furthermore, according to these authors, ground-finches are an appropriate model to study forces involved in geographic variation and local adaptation, but not to demonstrate the workings of speciation because in their view speciation in the group has not occurred. Instead, incipient speciation has been repeatedly stalled or reversed owing to shifting conditions affecting the strength and direction of natural selection and to ongoing gene flow, a situation they wittily referred to as “Sisyphean” evolution (McKay and Zink 2015). Because of its originality in challenging “entrenched orthodoxy regarding speciation in Darwin’s Finches”, the study by McKay and Zink (2015) was duly recognized with an award by a major ornithological organization (Cooper Ornithological Society 2016).

A central premise of the arguments by McKay and Zink (2015) was their assertion that phenotypic discontinuities do not exist among recognized species of ground-finches (contra Lack 1947; Grant et al. 1985). Although they rightly noted that “the real test of species limits is determining the extent to which specimens form multiple morphological clusters when *a priori* specimen identifications are ignored”, McKay and Zink (2015) did not formally conduct such a test. Instead, their approach illustrates three problematic issues in analyses of phenotypic data for species delimitation. In the next section we describe these issues and outline possible solutions afforded by statistical tools directly related to the basic evolutionary model underlying the use of phenotypic data in species delimitation. We then implement these solutions in a reanalysis of the morphological data on *Geospiza* ground-finches to revisit the question of whether morphological evidence supports the hypothesis that there are several species in the group.

## Three Frequent Issues in Analyses of Phenotypic Data for Species Delimitation

### 1) Graphical analyses may convey little information on phenotype frequencies crucial to assess evidence for multiple species

Many species delimitation studies rely on visual inspection of bivariate (rarely trivariate) scatter plots of phenotypic space to detect discontinuities and thus define phenotypic groups (e.g., Fig. 1 in McKay and Zink 2015). These scatter plots may offer only limited insight into the structure of character variation because visual cluttering and record overplotting hinder perception of phenotype frequencies crucial to identify groups (McLachlan 2004). We illustrate this problem with a hypothetical example in which specimens from a given locality seem to reveal no phenotypic discontinuities, with intermediate phenotypes across the range of variation (Fig. 1a); accordingly, a univariate scatter plot fails to reveal evidence of distinct phenotypic groups (Fig. 1b). The problem with scatter plots concealing crucial information (also common in two- and three-dimensional scatter plots) is revealed by a histogram of phenotype frequencies employing the same data, which reveals two distinct normal distributions (Fig. 1c). Following the model for species delimitation based on continuous phenotypic characters described above, this histogram suggests the existence of two species.

**Figure 1.**
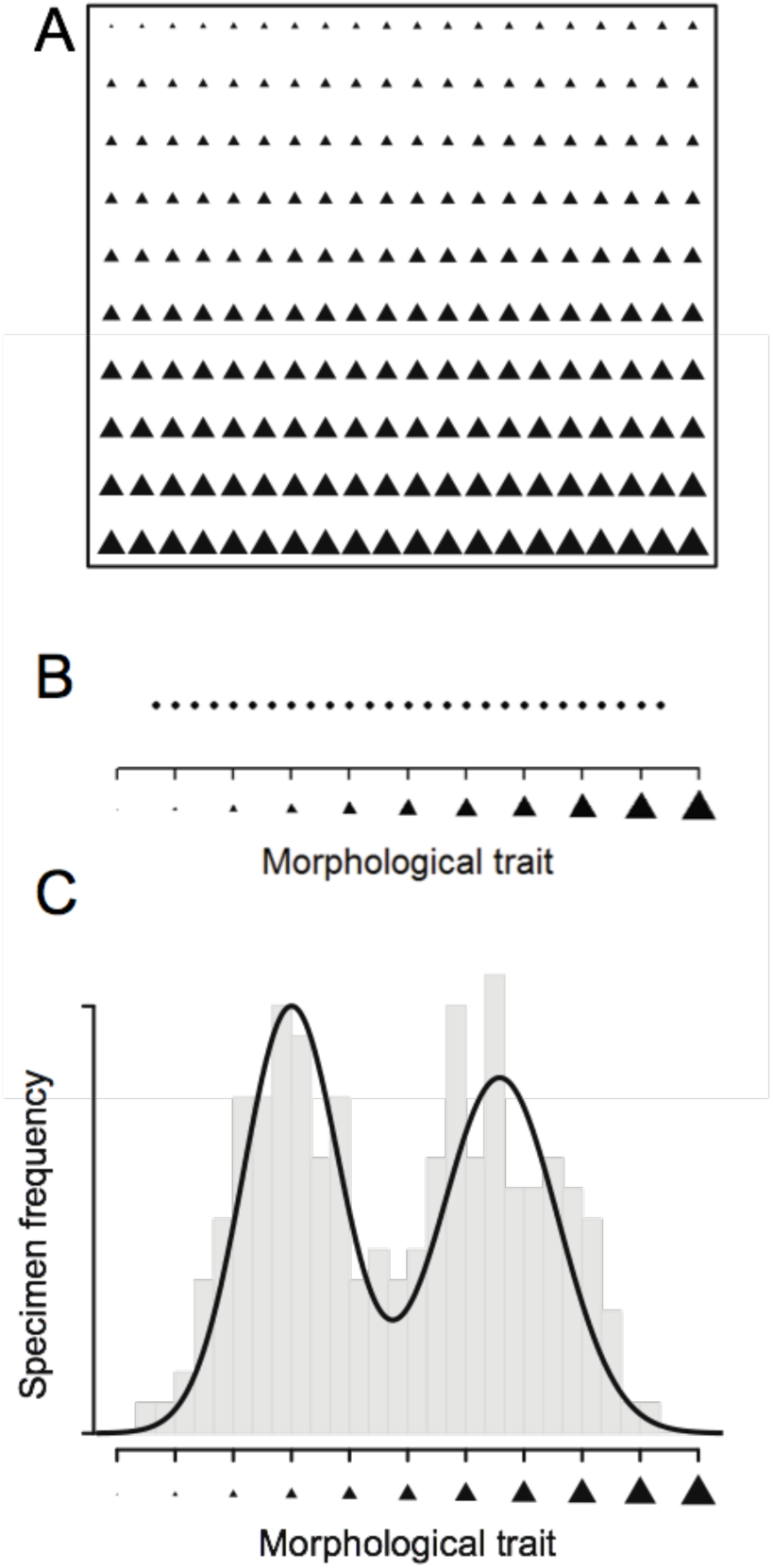
Visual inspection of phenotypic data may yield limited insight regarding species limits. (A) Sample of 200 museum specimens (triangles) arranged according to a morphological phenotype (triangle size), from small in the upper left to large in the lower right. The specimens appear to form a smooth gradient with no morphological gaps. (B) Plot of specimen measurements along a single continuous axis representing the size of the morphological trait in A. At the resolution of the measurements, extreme values in the sample seem to be gradually connected by intermediate phenotypes throughout. Thus, there seem to be no obvious morphological gap, suggesting the specimens correspond to a single variable species. (C) Two distinct normal distributions are revealed by examining the frequency (gray bars) of specimen phenotypes in the sample, suggesting that specimens may correspond to two species. In fact, the sample was drawn from a mixture of two normal distributions (continuous black lines).

Graphical analysis of phenotype frequencies (e.g., Fig. 1c) may be effective to detect groups when few characters are relevant (but see McLachlan and Peel 2000, page 9). However, it may be difficult to detect distinct normal distributions in phenotypic spaces defined by more than two dimensions, where complex covariance structures are likely (McLachlan 2004). Moreover, detection of phenotypic groups exclusively based on graphical analysis is potentially highly subjective and difficult to replicate, or, as stated by Karl Pearson (1894) over a century ago: “To throw the solution on the judgment of the eye in examining the graphical results is, I feel certain, quite futile”. Therefore, graphical analysis of phenotype frequencies is a useful but limited tool for species delimitation.

Recent statistical developments allow systematists to go beyond graphical analysis by using normal mixture models (NMMs, McLachlan and Peel 2000) as a formal approach to test for the existence of distinct species based on multivariate phenotypic data (Ezard et al. 2010; Guillot et al. 2012; Edwards and Knowles 2014; Kleindorfer et al. 2014). These models conceptualize phenotypic variation as a combination (i.e., a mixture) of distinct normal distributions; a mixture may include one or more distinct normal distributions, representing the hypothesis of one or more species, respectively. The parameters of a NMM specifying a particular hypothesis include the means and variance-covariance matrices describing the Gaussian phenotypic distribution of each species. These parameters can be estimated using maximum likelihood from data on phenotypic measurements, without *a priori* knowledge of species limits, employing the expectation–maximization algorithm (McLachlan and Krishnan 2008). Comparison of empirical support among models representing different hypotheses is often based on the Bayesian Information Criterion (BIC; Schwarz 1978), which evaluates the likelihood of each model while adjusting for model complexity (Fraley and Raftery 2002).

### 2) Reduction of dimensionality via PCA may exclude important characters for species delimitation

Species delimitation studies often begin analyses by reducing the dimensionality of phenotypic space, typically via principal component analysis (PCA) or related procedures (McLachlan 2004; Ezard et al. 2010), and then focus attention on few principal components accounting for most of the variation in the data. For example, McKay and Zink (2015) focused on three principal components explaining 99% of the variation in six morphological characters of *Geospiza* ground-finches (see their Fig. 1). This use of PCA and related procedures in taxonomy was suggested decades ago (Sneath and Sokal 1973) and is still prescribed nowadays (e.g., Ezard et al. 2010). However, there is no reason to believe that principal components accounting for most of the variation in a dataset are most useful for group discrimination (Chang 1983).

To illustrate the problem of reducing dimensionality to the principal components accounting for most of the variation, we use a hypothetical example based on two phenotypically distinct species, each represented by a bivariate normal distribution (Fig. 2a). The first principal component of the mixture of these two distributions explains >99% of the variation and, yet, it is useless to distinguish the two species (Fig. 2b). In contrast, the second principal component accounts for <1% of the variation and perfectly discriminates species (Fig. 2c). This example is bivariate for simplicity, but the statistical principle applies to mixtures of two normal distributions in any number of dimensions (Chang 1983). We stress that the problem at hand is not rotation of the data using PCA or related procedures, because such rotation may serve a number of useful purposes; rather, the problem is employing the amount of phenotypic variance explained by each principal component as a criterion to judge its usefulness to distinguish phenotypic groups (Chang 1983).

**Figure 2.**
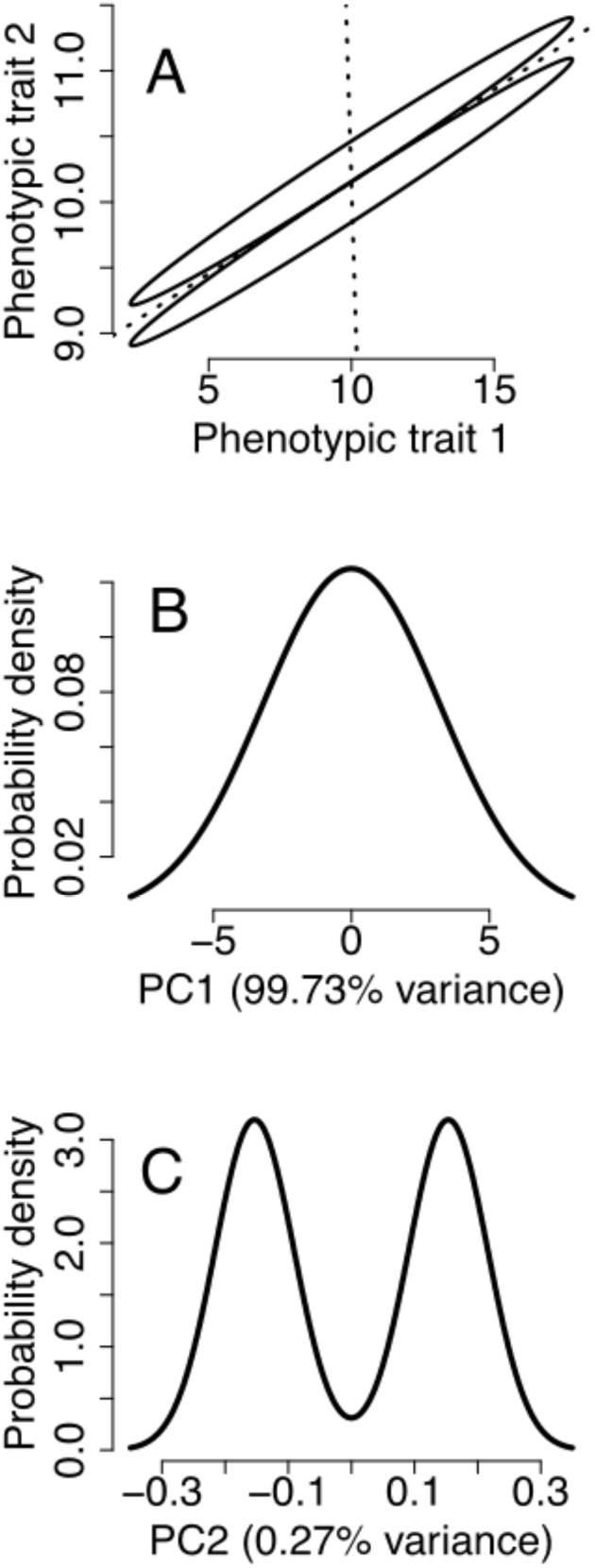
The ability of principal components to discriminate species is not necessarily proportional to the total phenotypic variance they explain. (A) Hypothetical example of two distinct species in the space defined by two phenotypic traits. Each species is described by a bivariate normal distribution, shown as ellipses covering 95% of the individuals of each species. Dotted lines represent the two principal component axes of the normal mixture of the two species. Note that the two principal components are orthogonal, forming a right angle that may not be apparent due to the magnification of the ordinate relative to the abscissa. (B) Probability density of individuals of the two species along the first principal component (PC1). This axis is useless to discriminate the phenotypic distributions of the two species, despite the fact that it explains 99.73% of the variance. (C) Probability density of individuals of the two species along the second principal component (PC2). The phenotypes of the two species can be readily distinguished along this axis, even though it explains only 0.27% of the variance. Because systematists often discard phenotypic axes accounting for small fractions of the total variance, they may miss crucial phenotypic evidence for species limits.

Although alternatives to PCA and related approaches for dimensionality reduction should be regularly considered in analysis aiming to detect groups in multivariate space (McLachlan and Peel 2000; McLachlan 2004), they are rarely implemented in species delimitation studies. For example, one may reduce dimensionality based on *a priori* considerations about which set of characters may be best to diagnose particular species. In particular, when *a priori* information about specific traits separating species is available (e.g. original species descriptions), one should favor analyzing variation in such traits; far from being circular (McKay and Zink 2015), it is only natural that one should critically examine evidence for species limits precisely in the dimensions in which such limits are hypothesized to exist (see also Remsen 2010; Patten and Remsen 2017). Alternatively, one may use methods that aim to find the set of variables (phenotypic traits) that best discriminates groups in a NMM, with no *a priori* information about groups (Raftery and Dean 2006; Maugis et al. 2009a; Maugis et al. 2009b).

### 3) Differences in central tendency are not evidence of distinct phenotypic distributions

Distinct normal distributions in quantitative characters constitute evidence for the existence of distinct species, but differences in central tendency between groups of individuals do not. This issue has been pointed out previously (e.g., Mayr et al. 1953; Luckow 1995; Patten and Unitt 2002), but seems to be ignored when statistical procedures to investigate differences in central tendency (e.g., t-tests, analysis of variance, Cohen’s d) are advanced as potentially valid tools to evaluate species boundaries (e.g., Simpson 1951; Henderson 2006; Tobias et al. 2010). McKay and Zink (2015, page 695 and their Fig. 2) have done as much by suggesting that statistical differences in average phenotypes between allopatric island populations of ground-finches could be equated to distinct morphological groups which, in turn, would have to be recognized as species.

Because this issue seems to commonly afflict assessments of species limits between allopatric forms (Tobias et al. 2010; McKay and Zink 2015), we illustrate it with a hypothetical example of two allopatric populations. Grouping specimens from these populations according to collection localities (e.g., two islands) reveals statistically significant differences in the central tendency of phenotypes (Fig. 3a). Yet, there is no evidence that phenotypic variation across specimens from the two populations is best described by more than a single normal distribution (Fig. 3b). Therefore, there is no evidence for more than one species in the sample of specimens regardless of differences in average phenotypes. This illustration focuses on groups of specimens defined by geography (i.e., allopatric populations), but the issue may affect comparisons involving groups of specimens defined by time (i.e., allochronic populations; Simpson 1951) or by any other criterion.

**Figure 3.**
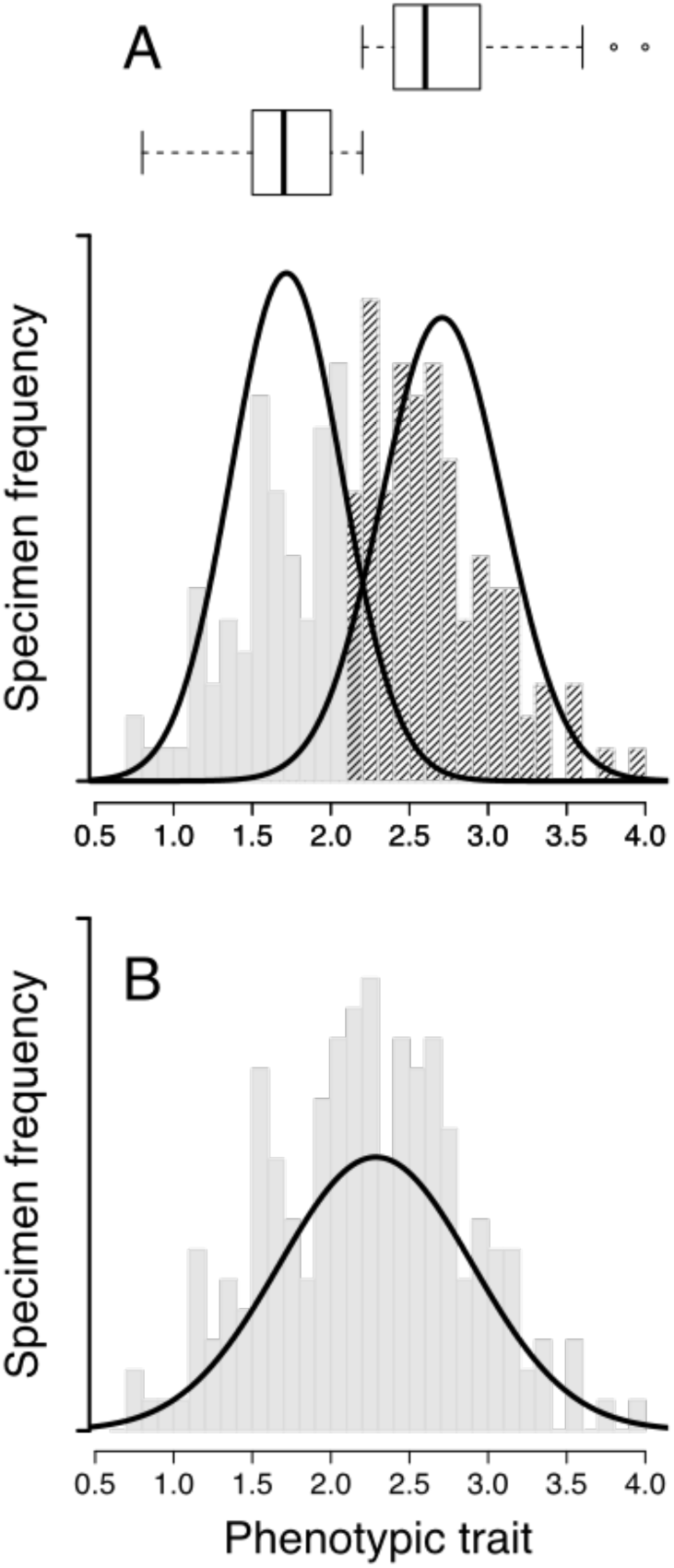
Differences in central tendency are not evidence of distinct phenotypic distributions. (A) Specimens from two allopatric populations (gray and striped histogram bars) differ markedly in the central tendency of a phenotypic trait (abscissa), as described by normal probability density functions (continuous black lines) and boxplots on top. The boxplots show the median (solid thick line), the interquartile range (box), whiskers extending to the most extreme values within 1.5 × interquartile ranges from the box, and outliers. The difference between means (1.717 and 2.707) is statistically significant (0.99, 95% CI: 0.89 - 1.09, t-test p-value < 2×10^−16^). Cohen’s d = 2.66, which is generally regarded as a large effect size. Note that the phenotypic ranges of the two populations do not overlap. (B) Despite the difference in central tendency and absence of phenotypic overlap, a single normal distribution (continuous black line) describes phenotypic variation across all specimens (gray bars) better than two normal distributions. In particular, empirical support for a normal mixture model assuming two distinct normal distributions (A) is substantially lower than that for a model specifying a single normal distribution (B): a difference of 10 in Bayesian Information Criterion (Schwarz 1978, Fraley and Raftery 2002). Thus, in light of the basic model for species delimitation based on quantitative phenotypic characters, there are no grounds to suggest that the specimens represent two distinct species despite marked differences in central tendency.

The solution to the problem is simple: do not treat phenotypic differences in central tendency as evidence for the existence of distinct phenotypic groups and, therefore, distinct species. No matter how statistically significant, even very large effect sizes are not germane in light of the basic model for species delimitation based on quantitative phenotypic characters. In light of this model, the focus of analysis should be on determining the number of normal distributions needed to describe phenotypic variation among specimens, as well as on estimating the parameters of those distributions (e.g., means and variance-covariance matrices). Indeed, strong evidence may exist for more than one distinct normal distribution in the absence of differences in central tendency (Hennig 2010), suggesting biologically meaningful differences between species in the variance of phenotypic traits (Supplementary Material, Appendix 1). As explained above, NMMs are a useful tool to test for distinct normal distributions.

## Are There Phenotypically Distinct Groups of Ground-Finches?

We examined phenotypic variation among *Geospiza* ground-finches by analyzing data from six morphological measurements of museum specimens (wing length, tail length, tarsus length, bill length, bill width, and bill depth) taken on adult males by H. S. Swarth for his monographic revision of the birds of the Galapagos (Swarth 1931). These were the same data employed by McKay and Zink (2015), which we use here with permission from the California Academy of Sciences; our sample sizes differ from those of the earlier study (501 vs. 486 male individuals) because we excluded a few individuals that were duplicated in the original dataset. The data we employed and the R code used to conduct the analyses described below are available as supplementary material (Appendices 2 and 3, respectively).

We asked how many distinct groups of ground-finches exist in the Galapagos using morphological data from specimens collected across the archipelago (total 18 islands). To define the morphological space for this analysis, we followed McKay and Zink (2015) and used PCA on the covariance matrix of log-transformed data. Rather than examining evidence for species limits using only the first three principal components accounting for >99% of the variation (McKay and Zink 2015), we used the R package clustvarsel (Scrucca and Raftery 2004) to reduce the dimensionality of the data by selecting the set of principal components most useful for group discrimination in NMMs, without *a priori* information about groups (Raftery and Dean 2006; Maugis et al. 2009a; Maugis et al. 2009b). We used the R package mclust 5.0 (Scrucca et al. 2016) to fit a wide range of NMMs. At one extreme, NMMs assuming one morphological group represented the Sisyphean evolution hypothesis that there is a single species of ground-finch (McKay and Zink 2015). Toward the opposite end, NMMs assuming up to 30 distinct morphological groups represented hypotheses alluded to by McKay and Zink (2015) when they suggested ground-finches may comprise “dozens of cluster species”, or “1 or 6 or 30 species” (p. 695). We also fitted NMMs specifying the six (Lack 1947) or nine (Lamichhaney et al. 2015; Remsen et al. 2017) species recognized by alternative taxonomic treatments of *Geospiza*, using the original specimen identifications in Swarth’s data updated to reflect changes in nomenclature. We used the Bayesian Information Criterion (BIC; Schwarz 1978) to measure empirical support for different NMMs (Fraley and Raftery 2002) and thereby explicitly evaluated the hypothesis that there is only one species of ground-finch (McKay and Zink 2015) relative to hypotheses that there are several species in the group (Lack 1947; Lamichhaney et al. 2015; Remsen et al. 2017).

We found the first four principal components to be most useful for group discrimination; NMMs ignoring the fourth principal component, although it explained only 0.006% of the morphological variance, had substantially less empirical support (ΔBIC ≥ 55) than those including it. Therefore, in contrast to McKay and Zink (2015), we did not discard the fourth principal component for analysis. The models specifying seven and eight distinct morphological groups of ground-finches received the strongest support ΔBIC ≤ 1.26). Support for all other models was considerably lower (ΔBIC in all cases >20; Fig. 4). In turn, the model with the lowest support represented the Sisyphean evolution hypothesis proposing no distinct morphological groups of ground-finches (i.e., that there is a single group; McKay and Zink 2015), which had a 500 BIC difference to the second-worse model and > 821 BIC difference to the two best models. Relative to the best models, those specifying groupings consistent with taxonomy recognizing six or nine species were weakly supported (Fig. 4), considering differences in BIC scores greater than 6 are typically regarded as strong or very strong evidence against models with lower support (Kass and Raftery 1995). In sum, the data provided poor empirical support for the hypothesis that ground-finches consist of only one species (McKay and Zink 2015) and strongly supported hypotheses of several morphologically distinct groups (Fig. 5, Supplementary Fig. 1); however, those groups did not readily align with existing taxonomic treatments of *Geospiza* (Fig. 6, Supplementary Fig. 2).

**Figure 4.**
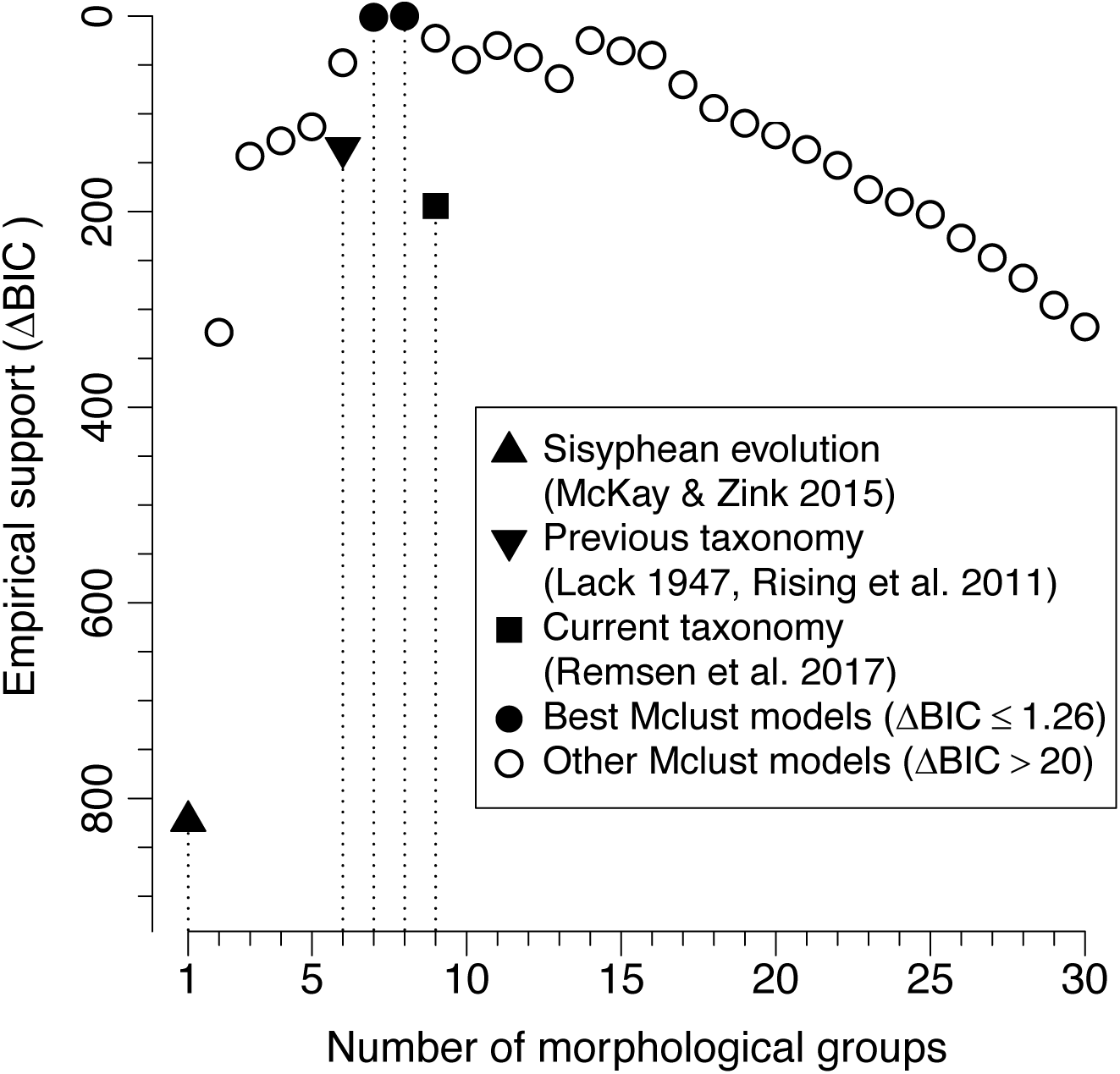
Analysis of morphological data strongly supported hypotheses that there are multiple distinct groups of *Geospiza* ground-finches. The plot shows the empirical support (ordinate) for normal mixture models assuming 1-30 distinct morphological groups (abscissa), and for the two models specifying groupings of specimens reflecting taxonomic treatments recognizing six species (Lack 1947, Rising et al. 2011) or nine species (Remsen et al. 2017). Empirical support was measured as difference in Bayesian Information Criterion relative to the best model (ΔBIC). The two models with highest empirical support assumed seven and eight distinct morphological groups. Empirical support for the model corresponding to the Sisyphean evolution hypothesis positing there is a single species of ground-finch (i.e. a single morphological group; McKay and Zink 2015) was negligible (ΔBIC > 820).

**Figure 5.**
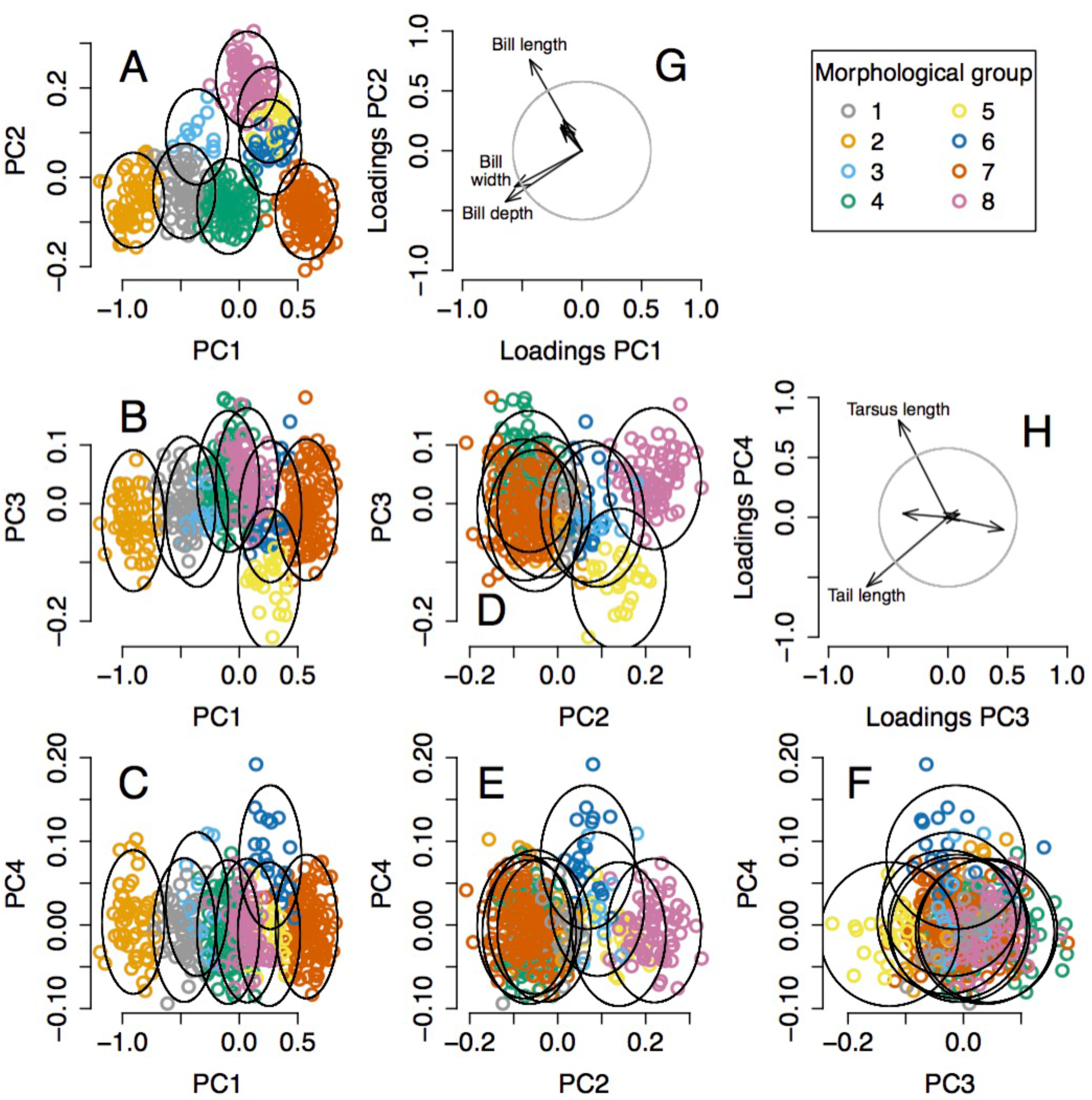
Eight morphological groups of *Geospiza* ground-finches identified by one of the best normal mixture models. Panels A-F show the groups in the space defined by the four principal components most useful for group discrimination (PC1-PC4). Colored symbols represent specimens assigned to different morphological groups and ellipses show 95% high density regions for normal distributions representing each morphological group. Arrows in G and H display the contribution of measured morphological traits to each principal component, gauged by the loadings of each trait on each principal component (i.e., elements of normalized eigenvectors). Circles show the length of arrows expected if all six traits contributed equally to bidimensional principal component spaces; arrows exceeding this expectation contribute most significantly and are labeled. PC1 and PC2 reflect general aspects of beak size and shape (G), with group 2 having long, deep and wide beaks, group 7 having short, shallow and narrow beaks, and morphological group 8 having long, shallow and narrow beaks. PC3 and PC4 reflect aspects of tail and tarsus length (H), with group 5 having a relatively long tail, and group 6 having a relatively long tarsus. PC4 is particularly useful to distinguish group 6 despite explaining only 0.6% of the total variance. The morphological distribution of groups in the other well supported NMM is fairly similar to the one shown here, the main difference being that group 3 is merged into groups 1 and 8 (Supplementary Fig. 1).

**Figure 6.**
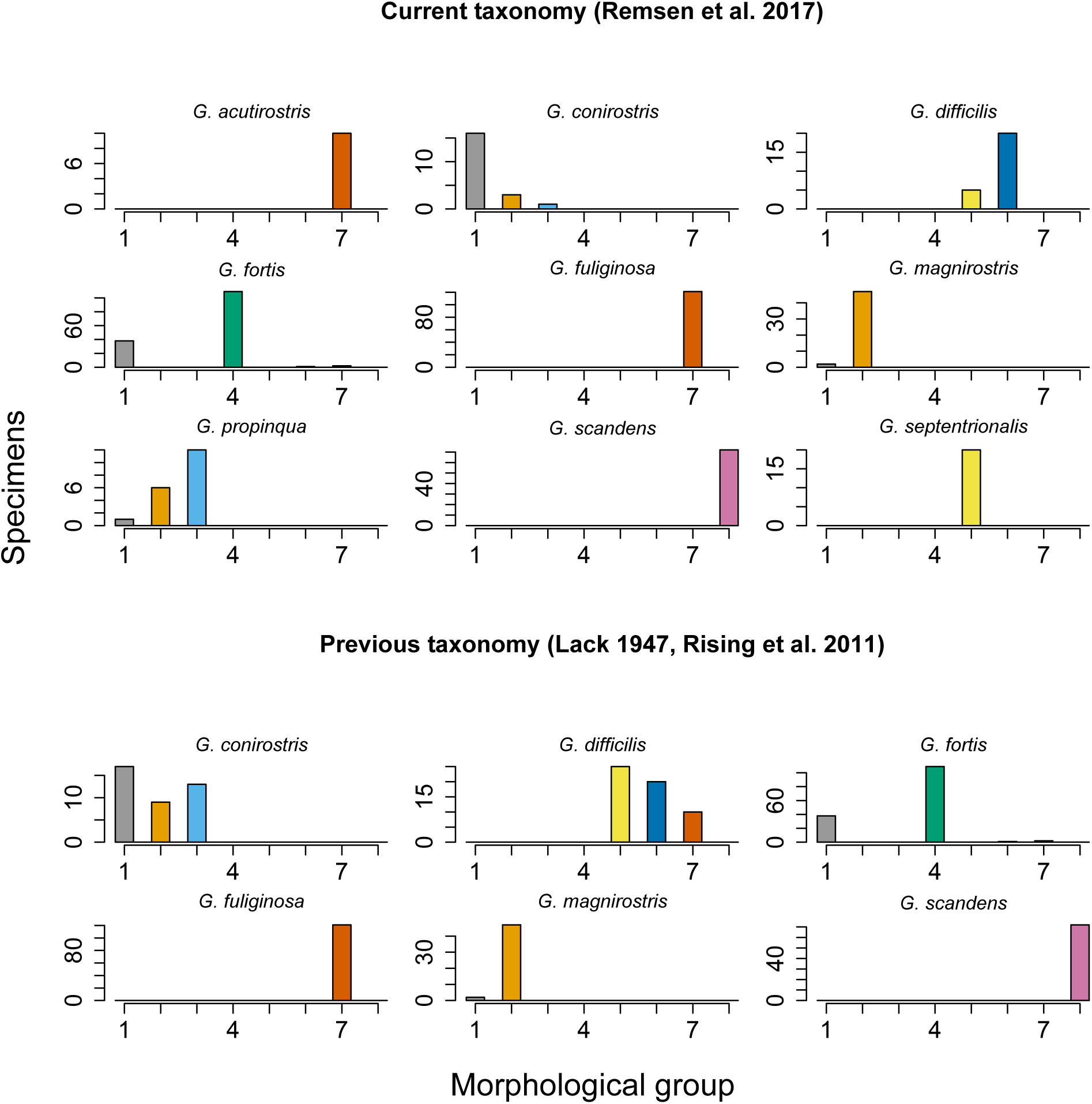
Eight morphological groups of *Geospiza* ground-finches in the Galapagos Archipelago identified by one of the best normal mixture models partially correspond to the nine species recognized by current taxonomy (Remsen et al. 2017) and to the six species recognized by previous taxonomy (Lack 1947, Rising et al. 2011). Each histogram shows, for each recognized species, the number of specimens assigned to each of the eight morphological groups. Groups are colored according to the scheme in Fig. 5.

Despite their comparatively low empirical support, models specifying six or nine morphological groups according to taxonomic treatments of *Geospiza* (Lack 1947; Remsen et al. 2017) were partially consistent with the best models (Fig. 6 and Supplementary Fig. 2). For example, in the best models, all specimens of two of the nine currently recognized species (*G. scandens, G. septentrionalis*) were assigned to two respective morphological groups which included few or no specimens of other species (Fig. 6; Supplementary Fig. 2). Discrepancies between our analysis and current taxonomy were most evident in cases such as those of (1) *G. propinqua, G. conirostris* and *G. fortis*, which were assigned to three, three (or two) and four morphological groups, respectively, or (2) *G. fuliginosa* and *G. acutirostris*, in which all specimens were assigned to the same group to the exclusion of nearly all specimens of other species. In addition, some morphological groups included specimens of multiple species (e.g., morphological group 1 contained specimens identified as *G. propinqua, G. fortis, G. magnirostris*, and *G. conirostris*). Part of the lack of agreement between the morphological groups we detected and groups recognized by taxonomy may be accounted for by considering that species may be told apart by phenotypic characters different from those we considered. For example, *G. fuliginosa* and *G. acutirostris* are indistinguishable in our analysis, but are distinct given subtle differences in bill profile and marked differences in songs (Grant and Grant 2008). Likewise, some of the discrepancies with current taxonomy (Remsen et al. 2017) involved cases in which species delimitation was not based on morphology, but rather resulted from recent genomic analyses revealing that phenotypically similar populations are distantly related (Lamichhaney et al. 2015). This likely explains why our analysis did not fully discriminate some species pairs in the morphological space we examined (*G. conirostris* vs. G. *propinqua, G. difficils* vs. G. *septentrionalis*), although they may be more distinct in other phenotypic spaces including bill profile and song as well as behavior (Grant et al. 2000; Grant and Grant 2002). Also, we assumed that specimen identifications in the data set we analyzed were faultless; thus, part of the apparent mismatch between morphological groups detected in our analyses and taxonomy may reflect identification errors. Evaluating this possibility would require detailed examinations of individual specimens beyond the scope of our work.

Geographic context is an important consideration in assessments of species limits using phenotypic traits. Under a wide range of species definitions (sensu de Queiroz 1998), distinct phenotypic groups among sympatric individuals are readily accepted as evidence for the existence of distinct species (Mayr 1992; Mallet 2008), whereas distinct phenotypic groups corresponding to non-sympatric populations may be less readily accepted as evidence of distinct species because they may reflect within-species differentiation due to geographic isolation or local adaptation (Zapata and Jiménez 2012). The morphological groups of ground-finches we detected (Fig. 5, Supplementary Fig. 1) cannot be interpreted to reflect within-species, among-island variation because these groups occurred on multiple islands and were sympatric with other groups; all of the morphological groups identified in the best NMMs were widely distributed across the Galapagos Archipelago (median = 8.5 or 9.0, range 3-14 islands per group; Table 1 and Supplementary Table 1) and most islands harbored several groups (up to 6 in Santiago and 7 in Santa Cruz; Fig. 7 and Supplementary Figure 3). Importantly, almost all morphological groups co-occurred with each other in at least one island; the only exception was morphological group 3 in one of the models, which co-occurred with four out of the other seven groups (Table 1 and Supplementary Table 1). McKay and Zink (2015) indicated that different morphs of ground-finches exist within islands and argued that if such morphs were treated as species, then one would need to recognize dozens of species in the group; our analysis suggests this is not the case given the occurrence of all morphological groups in multiple islands.

**Table 1.** Number of islands in the Galapagos Archipelago where each of the eight morphological groups of *Geospiza* ground-finches identified by one of the best normal mixture models were found to occur (diagonal) and co-occur with other groups (off diagonal). All groups co-occurred with each other in at least one island, except for cases involving group 3, which did not co-occur with three other groups. Note, however, that group 3 was not recovered as distinct in the other best model, which identified only seven groups (Supplementary Table 1).

|  | Group 1 | Group 2 | Group 3 | Group 4 | Group 5 | Group 6 | Group 7 | Group 8 |
| --- | --- | --- | --- | --- | --- | --- | --- | --- |
| Group 1 | 8 | 4 | 2 | 4 | 2 | 1 | 6 | 4 |
| Group 2 |  | 9 | 2 | 4 | 4 | 3 | 7 | 5 |
| Group 3 |  |  | 3 | 0 | 1 | 0 | 1 | 0 |
| Group 4 |  |  |  | 11 | 2 | 3 | 10 | 7 |
| Group 5 |  |  |  |  | 4 | 2 | 2 | 2 |
| Group 6 |  |  |  |  |  | 3 | 3 | 3 |
| Group 7 |  |  |  |  |  |  | 14 | 3 |
| Group 8 |  |  |  |  |  |  |  | 9 |

**Figure 7.**
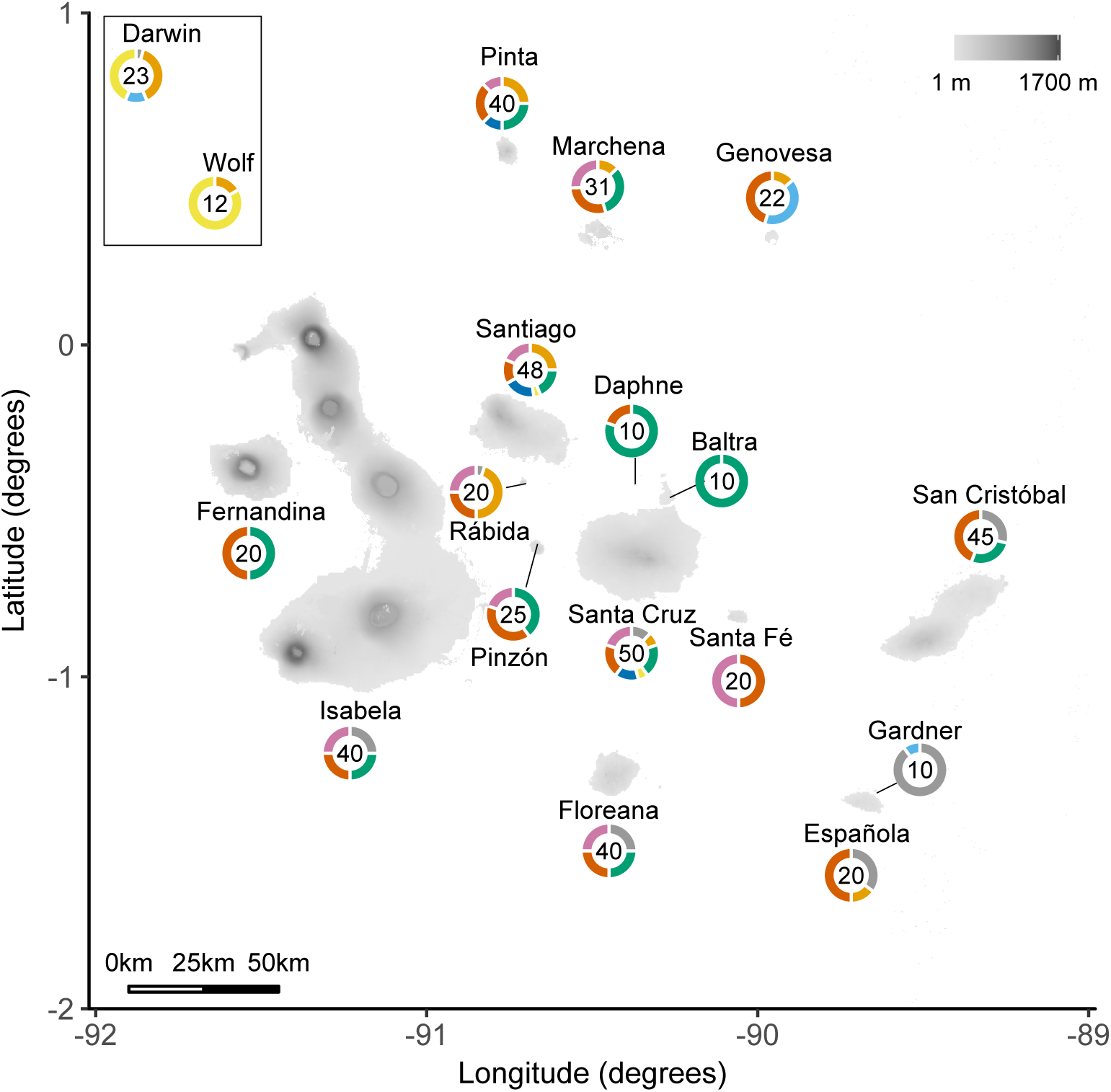
Eight distinct morphological groups of *Geospiza* ground-finches identified by one of the best normal mixture models have broad geographic distributions across the Galapagos Archipelago. For each island, numbers indicate individuals included in the analysis and ringplots depict the fraction of such individuals assigned to each morphological group following the color scheme in Fig. 5. The existence of distinct morphological groups in potential sympatry within islands (e.g., >4 groups in Santa Cruz, Santiago, and Pinta) suggests that such groups are unlikely to reflect within-species differentiation due to geographic isolation or local adaptation.

At this point we note that because the specimens we analyzed were collected several decades ago (Swarth 1931), they may not faithfully reflect patterns in morphological variation nor the geographic distributions of morphological groups in the present. This is because over the past century, ground-finch populations have experienced a few colonization and extinction events, changes in the degree of morphological differentiation among populations due to natural and human-mediated hybridization, and bouts of selection in shifting directions over multiple generations in association with environmental variation in space and time (Harris 1973; De León et al. 2011; Grant and Grant 2014). Thus, we refrain from additional discussions about species limits involving comparisons of historical morphological data with contemporary evidence (e.g., genomics; Lamichhaney et al. 2015). Nonetheless, our analyses serve to demonstrate that statistically distinct morphological groups of ground-finches existed in the past, and we strongly suspect they still exist in the present. Accordingly, we suggest that the burden of proof for systematists proposing to lump ground-finches into a single species based on morphological data is on showing that distinct groups do not longer exist.

## Conclusions; or, Atlantean Evolution in Darwin’S Finches

Our reanalysis of morphological data pointed strongly to the existence of several groups of phenotypically distinct *Geospiza* ground-finches based only on six linear morphological measurements. In addition, we found evidence of distinct phenotypes in geographic scenarios (i.e., sympatry within islands) where one should not expect them if populations had not achieved evolutionary independence. specifically, because the variation in quantitative morphological traits we examined is polygenic (Abzhanov et al. 2004; Abzhanov et al. 2006; Lamichhaney et al. 2015; Chaves et al. 2016; Lamichhaney et al. 2016) and not caused by differences in sex or age (we restricted analyses to adult males), the existence of distinct phenotypic groups in areas where populations come into contact implies there are likely several species of ground-finches. Therefore, we contend that ground-finches are not an example of sisyphean evolution (McKay and Zink 2015), a term that could well apply to other systems in nature (Seehausen 2006; Nosil et al. 2009; Rudman and Schluter 2016). Instead, evolutionary forces maintaining populations of ground-finches apart are likely in place, just as in Greek mythology Atlas prevents the merging of the Earth and the sky with his shoulders. Ground-finches thus likely represent an example of what one might call “Atlantean evolution”. One, of course, does not need a new term to refer to speciation, but thinking of Atlas brings to mind *atlas*, a collection of maps, which reminds one of the central role of geography in speciation and in the basic model underlying species delimitation based on phenotypic variation.

The question of exactly how many species of Darwin’s ground-finches are there remains open and requires further attention to morphology, including careful scrutiny of discrepancies between morphological variation and taxonomy (e.g., Fig. 6). In addition, morphological variation should be further examined in light of biological factors including additional phenotypic characters, ecological niches, mating behavior, population dynamics, and patterns of genetic and genomic variation among populations (Grant 1999; Huber et al. 2007; Grant and Grant 2008; Farrington et al. 2014; Grant and Grant 2014; Lamichhaney et al. 2015; McKay and Zink 2015). Fruitful discussions about species limits in the group would likely start by addressing some of the additional thought-provoking arguments advanced by McKay and Zink (2015) that we did not touch on and which are beyond the scope of our work (e.g., the extent to which morphological groups are stable lineages over time or the evidence for the existence of distinct gene pools). Any such discussions, however, as well as discussions over species delimitation in other organisms, should bear in mind that phenotypic evidence for species limits is best assessed using statistical approaches appropriately grounded on evolutionary models.

## OUTLOOK

We do not claim that the approaches used here to analyze phenotypic data for species delimitation are free of problems. Issues such as estimation of the number of groups in NMMs (McLachlan and Peel 2000; McLachlan and Rathnayake 2014) or how to select variables for NMM analyses of multidimensional datasets (Poon et al. 2013) are critical areas of active research in statistics in which progress remains to be made. Despite these issues, however, we argue that the statistical tools we used are appropriate because they are directly related to the basic evolutionary model underlying species delimitation using phenotypic data (Fisher 1918). Moreover, these tools allow systematists to go beyond fairly limited graphical analysis, and to break free from problems resulting from reduction of dimensionality using PCA or related approaches and from comparisons of measurements of central tendencies. The value of embracing approaches with a solid theoretical basis despite limitations in their implementation in systematics is clear considering other developments in the field in which theory predated robust methodologies that subsequently blossomed. Such developments include the use of statistical methods to study species limits among fossil populations (Newell 1956), the application of probabilistic models to infer phylogenetic trees (Felsenstein 1981), time-calibration of molecular phylogenies (Kishino and Hasegawa 1990), and the estimation of species trees from gene trees (Maddison 1997).

Practical approaches to fit NMMs without *a priori* information about species limits offer a fresh perspective in inferences available to systematists. In the absence of these tools, it seemed reasonable to argue that species limits should be based on fixed phenotypic differences because continuous variation could only be subdivided using subjective criteria (Cracraft 1989; Davis 1997). Accordingly, overlap of phenotypic ranges has been conventionally stressed as a criterion to suggest samples of individuals are conspecific (e.g., Simpson 1951; Davis and Heywood 1963; Zink 2002; McKay and Zink 2015).

However, overlap in phenotypic ranges under the framework offered by NMMs is not relevant for species delimitation because (1) one may find strong empirical support for models in which the phenotypic ranges of distinct normal distributions overlap (e.g., Fig. 1 and 5), and (2) absence of range overlap does not imply strong empirical support for models with more than a single species (e.g., Fig. 3). Critically, absence of range overlap need not imply a phenotypic gap (i.e., a phenotypic region with low frequency of individuals) because continuous phenotypic variation can be arbitrarily split into mutually exclusive parts regardless of phenotype frequencies. On the other hand, although true phenotypic gaps (along with multimodality in phenotypic distributions) are sufficient to suggest species boundaries (Zapata and Jiménez 2012; Mallet 2013), they are not necessary to demonstrate such boundaries exist because NMMs specifying more than one species may be strongly supported in the absence of phenotypic gaps. An example of support for more than one normal distribution in the absence of phenotypic gaps was provided at the inception of NMMs: Karl Pearson inferred two groups among specimens of the shore crab (*Carcinus maenas*) from the Bay of Naples, even though the mixture of the groups was not bimodal and therefore they were not separated by a gap (Pearson 1894). Moreover, Pearson examined the possibility of inferring the existence of groups with different phenotypic variances but identical phenotypic means, which are by definition not separated by a gap (Supplementary Material Appendix 1).

To conclude, we note that the criteria for species delimitation discussed above are relevant in the context of ideas about the reality of species. In particular, it has been argued that if the hypothesis that species are real entities in nature is correct, then biological diversity should be a patchwork of phenotypic clusters delineated by gaps (Coyne and Orr 2004; Barraclough and Humphreys 2015). This prediction, however, would not follow from the hypothesis that species are real if, as we argue, phenotypic gaps are not necessary criteria for species delimitation. In other words, species may be real entities in nature even if phenotypic gaps are not major elements structuring biological diversity. Because statistical approaches related to NMMs now allow systematists to make unprecedented formal inferences about the existence of species even if they overlap in phenotypic space, they constitute particularly useful tools to describe the structure of biological diversity, a necessary step to understand the evolutionary processes that generated it.

## Acknowledgements

We thank the California Academy of Sciences (Jack Dumbacher) for allowing us to use and publish morphological data from H. S. Swarth’s archive. Bailey McKay kindly provided tabulated data. We thank Peter Grant, Van Remsen, Elizabeth Spriggs, Peter Stevens, and members C.D. Cadena’s laboratory group for discussion and helpful comments on the manuscript.

## Supplementary Material

### Appendix 1.

Biological significance of differences between species in variance of phenotypic traits despite equal phenotypic means.

In analyses of phenotypic data, normal mixture models may reveal the existence of two or more normal distributions with different variances but equal means (Pearson 1894, McLachlan and Peel 2000, Hennig 2010). However, the biological significance of such phenotypic patterns might not be readily evident (Hennig 2010). Traditionally, systematists examining evidence for species limits have emphasized differences between species in phenotypic means, which are a necessary condition for the occurrence of phenotypic gaps (i.e., phenotypic spaces with low frequency of individuals) between species. Therefore, it is reasonable to ask why might differences between species in the variance of phenotypic traits be meaningful despite equal phenotypic means. Here we present a simple numerical example to show that, in the context of the Fisherian model for species delimitation based on phenotypic data we described, differences between species in phenotypic variance may be biologically meaningful and thus relevant for species delimitation.

The example considers diploid organisms in which a polygenic trait, *z*, is determined by *n* diallelic loci lacking dominance relationships or epistasis:

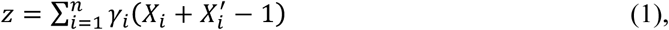
where *γ*_*i*_ is the allelic effect at locus *i*, and *X*_*i*_ and 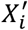 indicate which allele is present at locus *i* in two homologous chromosomes, respectively. Thus, *X*_*i*_ = 1 if the “+” allele is present in one chromosome and *X*_*i*_ = 0 otherwise. Likewise, 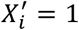 if the “+” allele is present in the other chromosome and 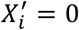 otherwise. Therefore, the (genetic) mean and variance of trait *z* are, respectively:

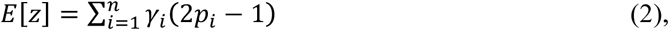

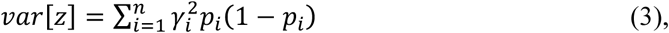
where *p*_*i*_ is the frequency of the “+” allele in locus *i* (de Vladar and Barton 2014). We assume eight unlinked loci (*n* = 8) of equal allelic effect (*γ*_*i*_ = 1 for all *i*). However, many examples with varying assumptions about number of loci and allelic effects are possible.

Imagine two sympatric species, A and B, for which the Fisherian model for species delimitation based on phenotypic data can be reasonably applied (see main text for assumptions of this model). Further imagine that allele frequencies at the eight unlinked loci determining trait *z* in each species are as follows:

| Locus | Species A | Species B |
| --- | --- | --- |
| 1 | $p_1 = 0.5$ | $p_1 = 1$ |
| 2 | $p_2 = 0.5$ | $p_2 = 0$ |
| 3 | $p_3 = 0.5$ | $p_3 = 1$ |
| 4 | $p_4 = 0.5$ | $p_4 = 0$ |
| 5 | $p_5 = 0.9$ | $p_5 = 0.9$ |
| 6 | $p_6 = 1$ | $p_6 = 1$ |
| 7 | $p_7 = 1$ | $p_7 = 1$ |
| 8 | $p_8 = 1$ | $p_8 = 1$ |

It can be seen, using equations 2 and 3, that the variance of trait *z* is higher in species A than in species B (1.09 and 0.09, respectively), despite a common trait mean (3.8 in both species). Thus, the differences in allele frequencies between species A and B are reflected in the variance of phenotypic trait *z*, and not in the mean of trait *z*. In the context of the Fisherian model for species delimitation based on phenotypic data, these differences in allele frequencies are biologically meaningful, because they would vanish after a few generations of random mating (Templeton 2006). In other words, variation in allele frequencies resulting in equal means but different variances between populations will only persist if such populations belong to different species; hence, differences in variances that one may detect employing normal mixture models are evidence supporting the hypothesis that there is more than one species in a sample of individuals.

### Appendix 2

Morphological measurements and geographic provenance of male specimens of *Geospiza* ground-finches employed in analyses are provided in a separate file **data.csv**. The data are from H. S. Swarth’s archive and were employed in this study and made available thanks to permission from the California Academy of Sciences.

### Appendix 3

R code employed to conduct analyses is provided in a separate file **Analysis_code.R**.

**Supplementary Figure 1.**
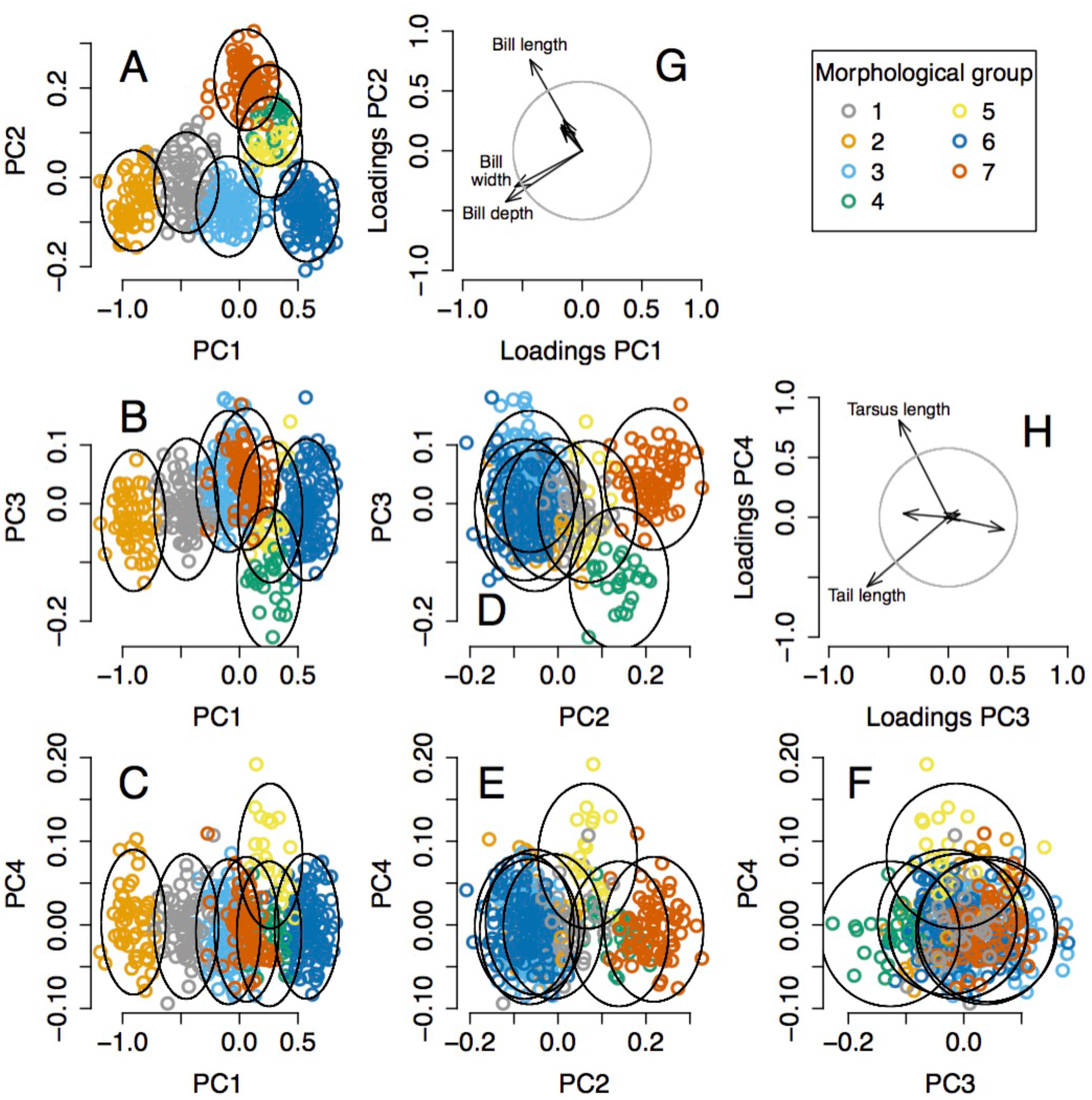
Seven morphological groups of *Geospiza* ground-finches identified by one of the best normal mixture models. Panels A-F show the groups in the space defined by the four principal components most useful for group discrimination (PC1-PC4). Colored symbols represent specimens assigned to different morphological groups and ellipses show 95% high density regions for normal distributions representing each morphological group. Arrows in G and H display the contribution of measured morphological traits to each principal component, gauged by the loadings of each trait on each principal component (i.e., elements of normalized eigenvectors). Circles show the length of arrows expected if all six traits contributed equally to bidimensional principal component spaces; arrows exceeding this expectation contribute most significantly and are labeled. PC1 and PC2 reflect general aspects of beak size and shape (G), with group 2 having long, deep and wide beaks, group 6 having short, shallow and narrow beaks, and group 7 having long, shallow and narrow beaks. PC3 and PC4 reflect aspects of tail and tarsus length (H), with group 4 having a relatively long tail, and group 5 having a relatively long tarsus. PC4 is particularly useful to distinguish group 5 despite explaining only 0.6% of the total variance. The morphological distribution of groups in the other well supported NMM is fairly similar to the one shown here, the main difference being that some individuals from groups 1 and 7 are placed together in an additional group (see Fig. 5 in main text).

**Supplementary Figure 2.**
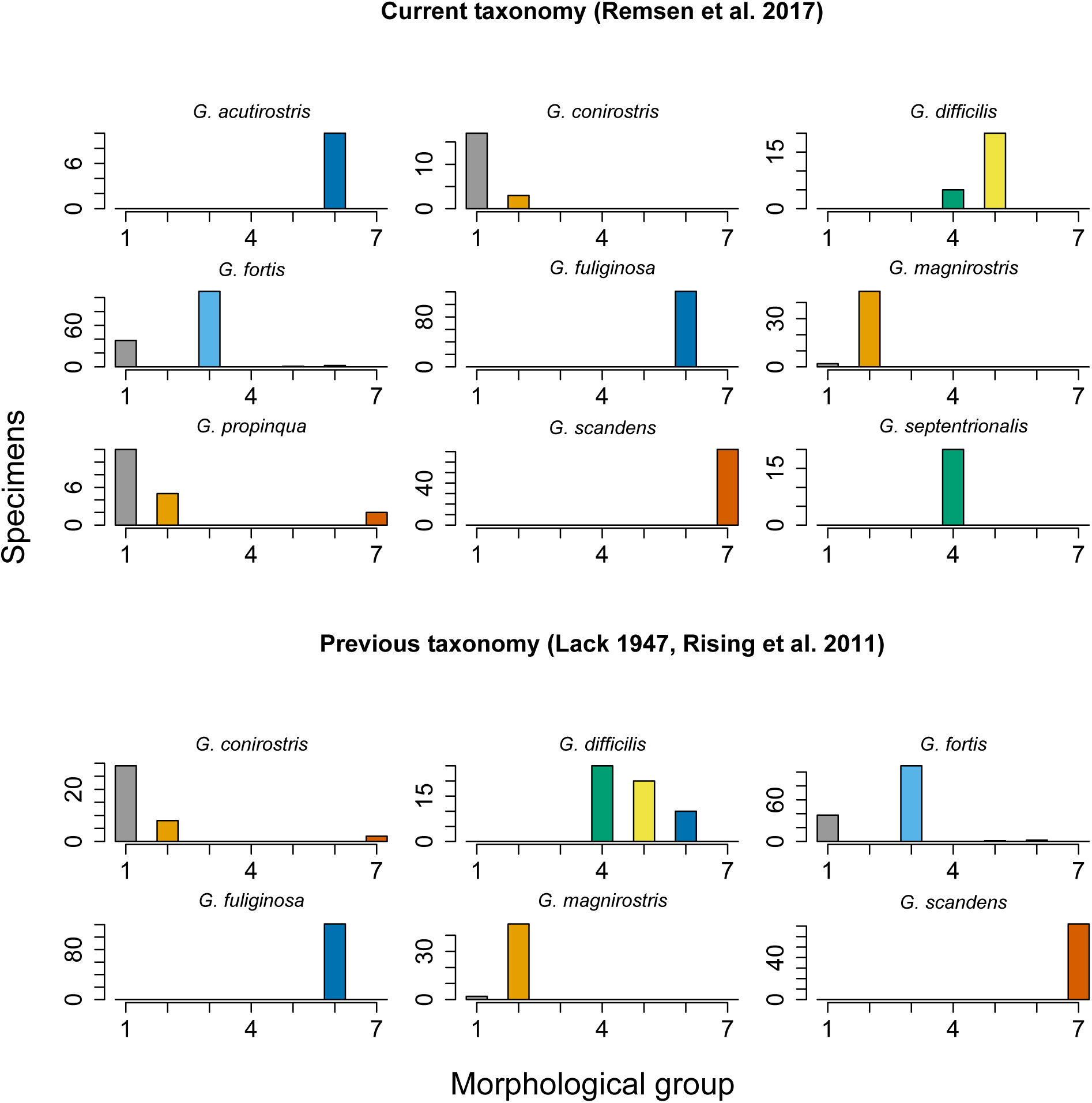
Seven morphological groups of *Geospiza* ground-finches in the Galapagos Archipelago identified by one of the best normal mixture models partially correspond to the nine species recognized by current taxonomy (Remsen et al. 2017) and to the six species recognized by previous taxonomy (Lack 1947, Rising et al. 2011). Each histogram shows, for each recognized species, the number of specimens assigned to each of the eight morphological groups. Groups are colored according to the scheme in Supplementary Fig. 1.

**Supplementary Figure 3.**
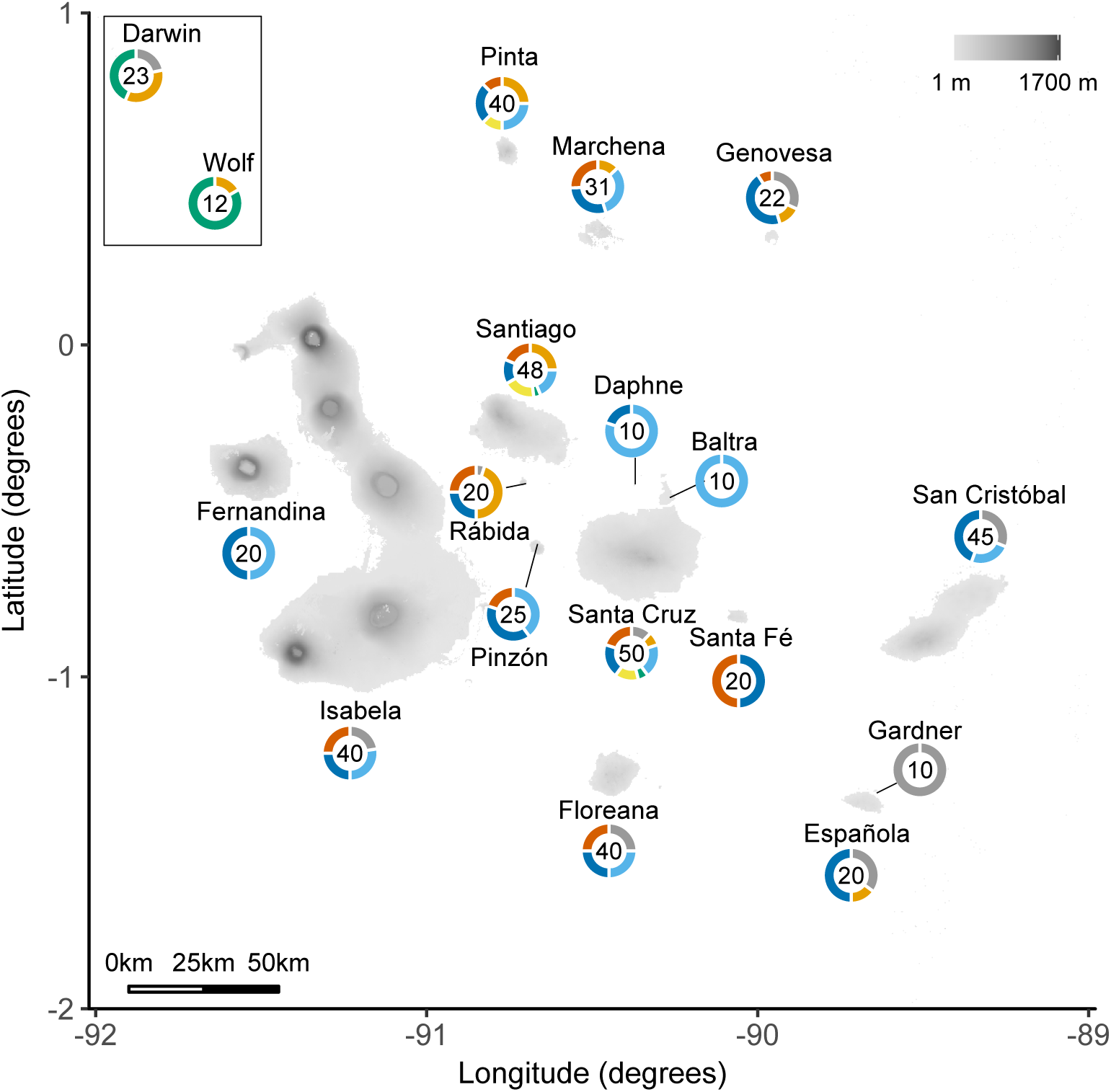
Seven distinct morphological groups of *Geospiza* ground-finches identified by one of the best normal mixture models have broad geographic distributions across the Galapagos Archipelago. For each island, numbers indicate individuals included in the analysis and ringplots depict the fraction of such individuals assigned to each of the eight morphological groups following the color scheme in Supplementary Fig. 1. The existence of distinct morphological groups in potential sympatry within islands (e.g., >4 groups in Santa Cruz, Santiago, and Pinta) suggests that such groups are unlikely to reflect within-species differentiation due to geographic isolation or local adaptation.

**Supplementary Table 1.** Number of islands in the Galapagos Archipelago 842 where each of the seven morphological groups of *Geospiza* ground-finches identified by one of the best NMMs were found to occur (diagonal) and co-occur with other groups (off diagonal). All groups co-occurred with each other in at least one island.

|  | Group 1 | Group 2 | Group 3 | Group 4 | Group 5 | Group 6 | Group 7 |
| --- | --- | --- | --- | --- | --- | --- | --- |
| Group 1 | 9 | 5 | 4 | 2 | 1 | 7 | 5 |
| Group 2 |  | 9 | 4 | 4 | 3 | 7 | 6 |
| Group 3 |  |  | 11 | 2 | 3 | 10 | 7 |
| Group 4 |  |  |  | 4 | 2 | 2 | 2 |
| Group 5 |  |  |  |  | 3 | 3 | 3 |
| Group 6 |  |  |  |  |  | 14 | 10 |
| Group 7 |  |  |  |  |  |  | 10 |

